# Camel nanobodies broadly neutralize SARS-CoV-2 variants

**DOI:** 10.1101/2021.10.27.465996

**Authors:** Jessica Hong, Hyung Joon Kwon, Raul Cachau, Catherine Z. Chen, Kevin John Butay, Zhijian Duan, Dan Li, Hua Ren, Tianyuzhou Liang, Jianghai Zhu, Venkata P. Dandey, Negin Martin, Dominic Esposito, Uriel Ortega-Rodriguez, Miao Xu, Mario J. Borgnia, Hang Xie, Mitchell Ho

## Abstract

With the emergence of SARS-CoV-2 variants, there is urgent need to develop broadly neutralizing antibodies. Here, we isolate two V_H_H nanobodies (7A3 and 8A2) from dromedary camels by phage display, which have high affinity for the receptor-binding domain (RBD) and broad neutralization activities against SARS-CoV-2 and its emerging variants. Cryo-EM complex structures reveal that 8A2 binds the RBD in its up mode and 7A3 inhibits receptor binding by uniquely targeting a highly conserved and deeply buried site in the spike regardless of the RBD conformational state. 7A3 at a dose of ≥5 mg/kg efficiently protects K18-hACE2 transgenic mice from the lethal challenge of B.1.351 or B.1.617.2, suggesting that the nanobody has promising therapeutic potentials to curb the COVID-19 surge with emerging SARS-CoV-2 variants.

**One-Sentence Summary:** Dromedary camel (*Camelus dromedarius*) V_H_H phage libraries were built for isolation of the nanobodies that broadly neutralize SARS-CoV-2 variants.

## Main text

Severe acute respiratory syndrome coronavirus 2 (SARS-CoV-2) is the etiologic agent of coronavirus disease 2019 (COVID-19) (*1, 2*) that enters human cells by binding its envelope anchored type I fusion protein (spike) to angiotensin-converting enzyme 2 (ACE2) (*3, 4*). SARS-CoV-2 spike is a trimer of S1/S2 heterodimers with three ACE2 receptor binding domains (RBD) attached to the distal end of the spike via a hinge region that allows conformational flexibility (*4*). In the all-down conformation, the RBDs are packed with their long axes contained in a plane perpendicular to the axis of symmetry of the trimer. Transition to the roughly perpendicular up conformation exposes the receptor-binding motif (RBM), located at the distal end of the RBD, which is sterically occluded in the down state. TMPR2 or cathepsin mediated proteolytic cleavage of an N-terminal fragment in each S2 monomer follows the binding of one or more RBMs to ACE2. This triggers a series of conformational changes in S2 that result in the fusion of the viral envelope to host cell membranes. The genetic payload of the virus is thus delivered into the cytoplasm. Neutralizing antibodies that target the spike, particularly its RBD, have been developed to treat COVID-19 by blocking the interaction with ACE2 and reducing SARS-CoV-2 infectivity (*5-9*). Most vaccines, including those mRNA-based, are designed to induce immunity against the spike or RBD (*10-14*). However, emerging SARS-CoV-2 variants such as B.1.1.7 (Alpha, UK), B.1.351 (Beta, South Africa), and P.1 (Gamma, Brazil) have exhibited increased resistance to neutralization by monoclonal antibodies or post-vaccination sera elicited by the COVID-19 vaccines (*15, 16*). Current monoclonal antibodies with Emergency Use Authorization for COVID-19 treatment partially (Casirivimab) or completely (Bamlanivimab) failed to inhibit the B.1.351 and P.1 variants. Similarly, these variants were less effectively inhibited by convalescent plasma and sera from individuals vaccinated with BNT162b2 (*15*). More recently, the new B.1.617.2 (Delta, India) variant became the prevailing strain in many countries (*17*). Thus, highly effective and broadly neutralizing antibody therapy is in urgent demand for COVID-19 patients.

Camelid V_H_H single domain antibodies (also known as nanobodies) can recognize protein cavities that are not accessible to conventional antibodies due to their small size and unique conformations (*18*). In the present study, we built novel camel V_H_H single domain antibody phage libraries from six dromedary camels (*Camelus dromedaries*). We used both the SARS-CoV-2 RBD and the stabilized spike ectodomain trimer protein as baits to conduct phage panning for nanobody screening. Among all the binders, NCI-CoV-7A3 (7A3), NCI-CoV-1B5 (1B5), NCI-CoV-8A2 (8A2), and NCI-CoV-2F7 (2F7) are potent ACE2 blockers. In addition, these nanobodies have potent neutralization activities against B.1.351 and B.1.1.7 variants and the original strain (Wuhan-Hu-1) as single agents. Interestingly, cross-competition assay and the cryo-electron microscopy (cryo-EM) structure of the spike trimer protein complex with these V_H_H nanobodies reveal two non-overlapping epitopes on the RBD for neutralization of SARS-CoV-2. The nanobody 8A2 directly interferes with the ACE2 binding on the RBD in its up mode. In contrast, 7A3 binds to a unique conserved epitope on the RBD regardless of its up or down conformational state. The 2-in-1 cocktail treatments, particularly 7A3 and 8A2, show more potent and broader activities against various variants in culture and protection of the mice infected with the B.1.351 variant. Interestingly, 7A3 retains its neutralization against the lethal challenge of the new B.1.617.2 variant in mice, indicating the nanobody has broad neutralization against all the major known variants.

## Results

### Isolation of camel nanobodies against SARS-CoV-2

To identify nanobodies against SARS-CoV-2, phage panning was carried out to screen RBD or the S protein using our new V_H_H single domain phage display libraries constructed from six camels, three males and three females, with ages ranging from 3 months to 20 years (Fig. 1A). Both SARS-CoV-2 RBD and the S trimer were used for phage panning. In total, 768 V_H_H phage clones were isolated; among them, 127 V_H_H clones have a high binding signal for the RBD. Out of the 127 clones, there were 29 unique V_H_H sequences (Fig. 1B). The top 6 V_H_H single domains that bind both the RBD and S protein of SARS-CoV-2 (including 1B5 which also binds the S1 subunit of SARS-CoV previously detected in 2003) were selected for further analysis. The sequence alignment of the top 6 shows that NCI-CoV-7A3 (7A3) and NCI-CoV-8A4 (8A4) are the V_H_Hs containing only two canonical cysteines, one before N-terminal to CDR1 and the other before CDR3 (Fig. S1). In contrast, the rest are the V_H_Hs, NCI-CoV-1B5 (1B5), NCI-CoV-8A2 (8A2), NCI-CoV-2F7 (2F7), and NCI-CoV-1H6 (1H6), having a total of 4 cysteines with two additional, non-canonical cysteines, one in CDR1 and the other in CDR3. The nanobodies with four cysteines are common in camels and sharks (*19*), suggesting they have antigen recognition mechanisms different from human and mouse IgG antibodies.

**Fig. 1.**
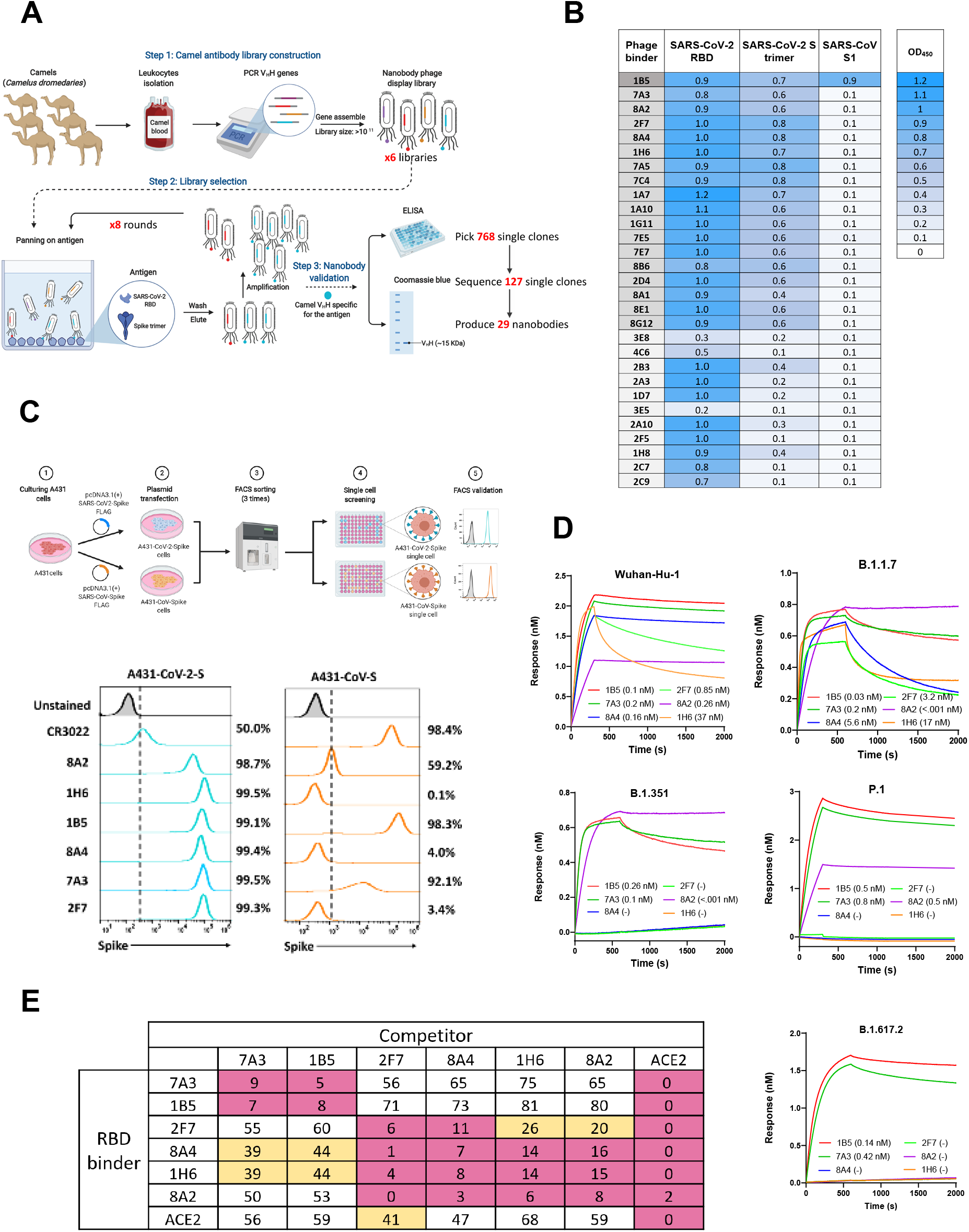
Isolation of high-affinity camel nanobodies against SARS-CoV-2. **(A)** Isolation of camel V_H_H nanobodies that bind the RBD phage display. **(B)** Camel V_H_H nanobodies against the S protein of SARS-CoV-2 or SARS-CoV. **(C)** Flow cytometry was performed to monitor the cross-reaction of nanobodies to the spike of both SARS-CoV-2 and SARS-CoV on cells. Outline of the experimental workflow for overexpression of SARS-CoV-2-spike or SARS-CoV-spike in the A431 human cell line. Both cell lines were stained with V_H_H nanobodies or CR3022 as a positive control. (**D)** Affinity binding (K_D_) of V_H_H-hFc antibodies against wild-type SARS-CoV-2 and mutants. The B.1.1.7 variant mutations include HV69-70 deletion, Y144 deletion, N501Y, A570D, D614G, P681H. The B.1.351 variant mutations include K417N, E484K, N501Y, D614G. The P.1. variant includes the E484K mutation. The B.1.617.2 variant includes the T19R, Δ (157-158), L452R, T478K, D614G, P681R, D950N mutations (**E)** Cross competition assay of each single domain antibody and ACE2 on Octet.

To validate the binding and cross-reactivity of the anti-RBD specific V_H_H nanobodies, we engineered two human cell lines, referred to as “A431-CoV-2-S” and “A431-CoV-S”, which overexpressed the S protein of either SARS-CoV-2 or SARS-CoV on the cell surface (Fig. 1C). CR3022 is used as a control antibody that is cross-reactive to both SARS-CoV-1 and SARS-CoV-2 RBD (*20*). All six anti-RBD nanobodies were able to bind A431-CoV-2-S cell lines with high affinity. Interestingly, 2 out of 6 anti-RBD V_H_Hs (1B5 and 7A3) showed cross-reactivity to A431-CoV-S with 8A2 having modest signals (Fig. 1C). These results indicated that the camel V_H_H nanobodies can bind the S protein of SARS-CoV-2 on the cell surface.

To measure the binding affinity against wild-type SARS-CoV-2 and emerging variants, B.1.1.7, B.1.351, and P.1, an Octet analysis was carried out (Fig. 1D, Table S1). For Wuhan-Hu-1 and B.1.1.7 variant binding, all six V_H_H-hFc fusion proteins exhibit subnanomolar binding affinity. Interestingly, for the B.1.351 and P.1 variants, only 8A2 (0.001 nM, 0.5 nM), 7A3 (0.1 nM, 0.8 nM) and 1B5 (0.26 nM, 0.5 nM) exhibited strong binding for the spike protein, whereas the rest V_H_Hs lost binding for these two variant spikes. More recently, for B.1.617.2 only 1B5 (0.14 nM) and 7A3 (0.42 nM) retained binding, whereas 8A2 lost the most of its binding to this new variant. Furthermore, a cross-competition assay showed two distinct epitope groups: 7A3 and 1B5 binds to a similar epitope, whereas the rest nanobodies (2F7, 8A4, 1H6, and 8A2) bind to a different epitope, indicating two non-overlapping epitopes on the RBD (Fig. 1E). Epitope mapping using an RBD-derived peptide array indicates that the nanobodies bind discontinuous conformation epitopes (Fig. S2). Additionally, we examined the inhibitory effect of V_H_H single domains against the RBD and the human ACE2 interaction (Fig. S3). Two V_H_Hs (1B5 and 8A2) were identified as the best ACE2 blockers among all V_H_Hs tested. 1B5 (IC_50_ = 3.2 nM) was the most potent ACE2 inhibitor followed by 8A2 (IC_50_ = 8 nM). Taken together, our data show there are two distinct epitopes on the RBD that can be recognized by our camel V_H_H nanobodies. One of them is recognized by 7A3 and 1B5 and conserved between SARS-CoV-2 and SARS-CoV; the other epitope is recognized by 8A2 and 2F7, unique for SARS-CoV-2.

### Camel nanobodies neutralize SARS-CoV-2 variants

To determine whether those nanobodies can neutralize SARS-CoV-2 entry, we tested them in pseudovirus neutralization assays in multiple systems using V_H_H-Fc fusions (Fig. 2A). Among the top 6 V_H_Hs, 8A2 is the best to inhibit the pseudovirus among six binders with an IC_50_ at 5 nM (Fig. 2B). We also used microneutralization assay to prescreen the top 6 V_H_Hs against the original strain and D614G variant and found that 8A2 showed the best neutralizing activity against early SARS-CoV-2 with or without D614G (Fig. S4-S5). This result was consistent with the pseudovirus neutralization result in Fig. 2B suggesting that 8A2 was more potent in blocking the entry of early SARS-CoV-2 than other 5 V_H_Hs. We then tested 2-in-1 and 3-in-1 combinations for all six binders in the pseudovirus infection assay. We found that 7A3+8A2 showed the best efficacy (IC_50_ of 1.6 nM) compared to the other combinations and single nanobodies (Fig. S4, Table S2). However, the combinations of 3-in-1 V_H_Hs did not further improve the efficacy (data not shown). During the process of characterizing these new V_H_H nanobodies, several variants of concern (VOCs) including B.1.1.7 (Alpha), B.1.351 (Beta), and P.1 (Gamma) emerged. To assess whether our nanobodies and combinations can neutralize these VOCs, pseudovirus for those mutants were produced and tested using 7A3, 8A2, 1B5, and 2F7 nanobodies. We found that the combination 7A3 and 8A2 showed the best efficacy in neutralizing the original virus and B.1.1.7, B.1.351, and P.1 variants with IC_50_ being 1 nM, 0.4 nM, 0.3 nM, and 0.2 nM, respectively (Fig. 2C and Table S2). Furthermore, fluorescence reporter-based pseudovirus data further validated luciferase-based pseudovirus data (Figs. S4). Together, our camel nanobodies exhibit potent neutralization activities against B.1.351, B.1.1.7, and P.1 variants, with the combination of 7A3 and 8A2 showing the best efficacy.

**Fig. 2.**
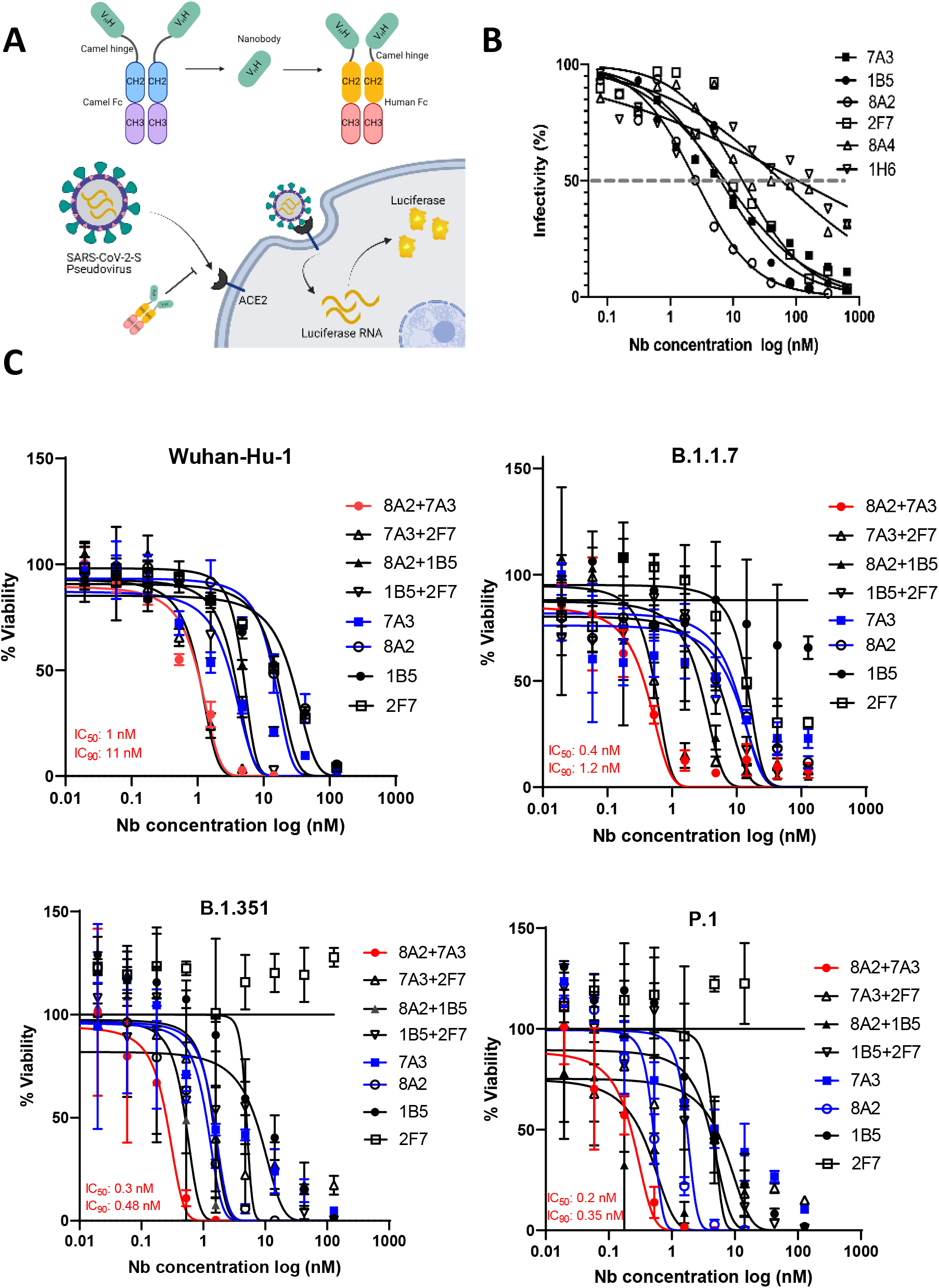
Nanobodies neutralize SARS-CoV-2 and the variants in pseudovirus assay. (A) Diagrams illustrating pseudovirus assay and V_H_H-hFc. (B) Camel V_H_H-hFc proteins inhibit SARS-CoV-2 pseudovirus infectivity to ACE2 expressing human cells by measuring luciferase expression. (C) Pseudovirus particle neutralization assay testing 2-in-1 combination and single nanobodies showing that 7A3+8A2 combination has the best neutralization activity.

To determine the best single domain antibody combination against the live virus, the combinations of 7A3, 2F7, 1B5, 8A2, and 8A4 V_H_H-hFc fusion proteins along with their single agents were tested using cytopathic effect assay along with the Wuhan-Hu-1 strain. Among them, 7A3+2F7 and 7A3+8A2 showed the best activity, with IC_50_ values being 16 nM and 20 nM, respectively (Fig. 3A, Table S3). To further test the combinations against variant viruses, three 2-in-1 combinations (8A2+7A3, 8A2+2F7, and 7A3+2F7) are tested in live virus-based microneutralization assay against the early SARS-CoV-2 strain bearing 614D, D614G variant, and VOCs including B.1.1.7, B.1.351, P.1, and B.1.617.2. In general, the combination of 8A2 and 7A3 showed potent neutralization activity across all the virus variants tested with IC_50_ values of 6 nM (D614G), 2 nM (B.1.1.7), 0.87 nM (B.1.351), 0.14 nM (P.1), and 27 nM (B.1.617.2) respectively (Fig. 3B, Table S3). While 8A2 lost neutralization against B.1.617.2, 7A3 alone exhibited better neutralization activity (19 nM) than the 2-in-1 combination against B.1.617.2 variant (Fig. 3B, Table S3). These results indicate that 7A3 has broad neutralization against all variants tested including B.1.617.2.

**Fig. 3.**
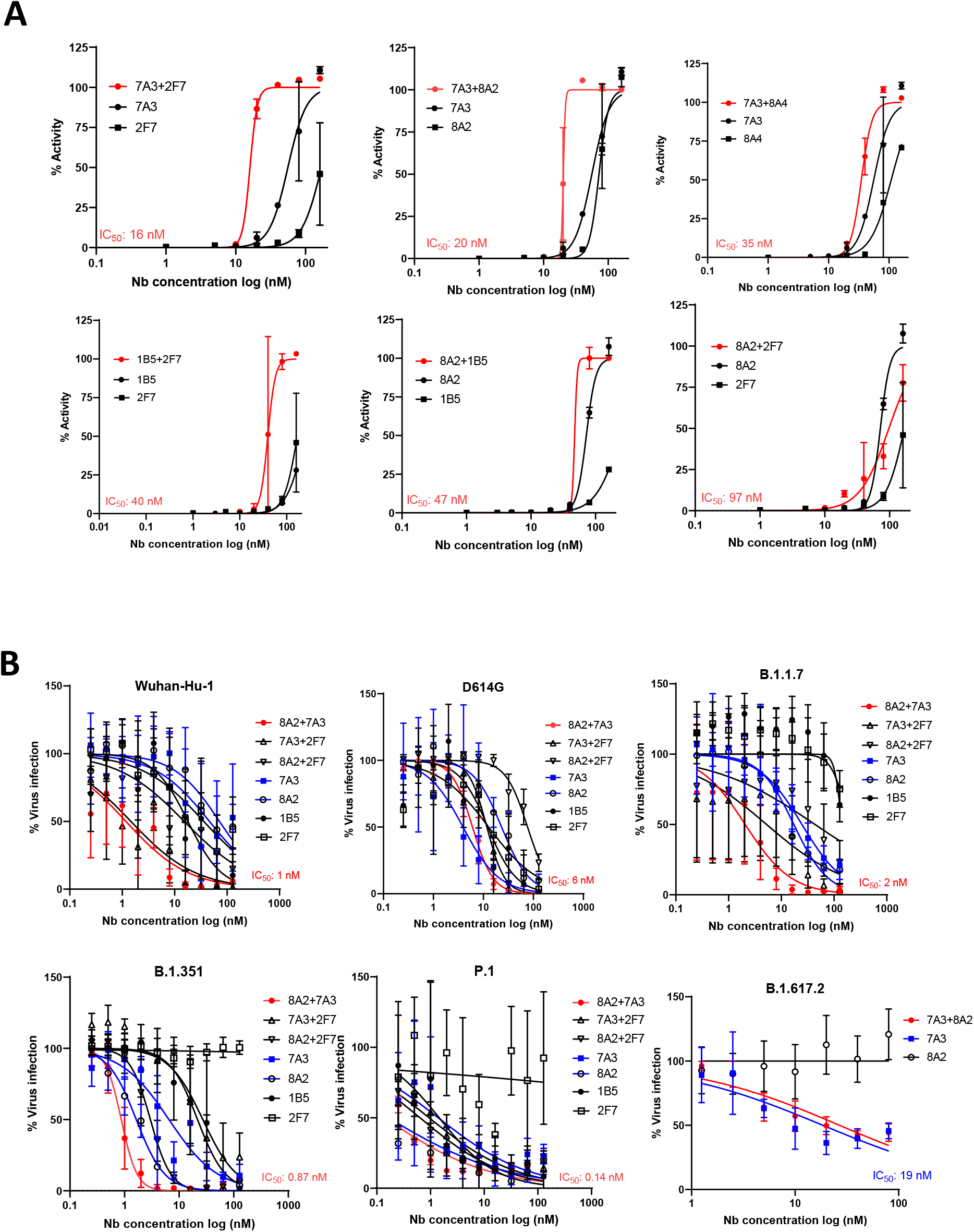
Neutralization of SARS-CoV-2 and its variants in live virus assay. **(A)** Live virus neutralization assay of nanobodies 7A3, 8A2, 2F7, 1B5, and 8A4 along with their 2-in-1 combination against Wuhan-Hu-1 strain. **(B)** Live variant virus assay using top three nanobody combinations of 8A2+7A3, 8A2+2F7, and 7A3+2F7 was conducted against SARS-CoV-2 (USA-WA1/2020), D614G variant, B.1.1.7 variant, B.1.351 variant, P.1 variant, and B.1.617.2 variant.

### Nanobodies protect K18-hACE2 transgenic mice from lethal B.1.351 and B.1.617.2 variant challenge

We then used the K18-hACE2 transgenic mouse model bearing human-like ACE2 to demonstrate in vivo efficacy of these nanobodies. K18-hACE2 mice infected with B.1.351 and B.1.617.2 variants had scattered foci of inflammation in lung and vasculitis and neuronal degeneration in brain as revealed by hematoxylin and eosin staining (Fig. 4A). The lethal infection of B.1.351 and B.1.617.2 resulted in 100% mortality and up to 30% body weight (BW) loss in virus only K18-hACE2 mice (Figs. 4B top panels). K18-hACE2 mice receiving 7A3 V_H_H-hFc fusion protein or the 2-in-1 cocktail of 7A3 and 8A2 at the dose of 5 mg/kg had 100% survival after the lethal B.1.351 infection with no BW drop within the 2-week observation period (Fig. 4B left panels). However, two out of four mice receiving 8A2 at 5 mg/kg died within 8 days after infection (Fig. 4B left panels). The surviving mice at 4 weeks after the lethal B1.351 infection had spike-specific IgG geometric mean titer (GMT) of 226 (the 2-in-1 cocktail group), 3620 (the 7A3 treatment), and 5120 (the 8A2 treatment), respectively (Fig. 4C). This indicated that the 2-in-1 cocktail or 7A3 alone at the dose of 5 mg/kg was more efficient than 8A2 alone to reduce viral load and protect K18-hACE2 mice from the lethal B.1.351 infection. When K18-hACE2 mice were subjected to a lethal dose of B.1.617.2, however, the protective efficiency of the 2-in-1 cocktail (5 mg/kg) was reduced to 50% and infected mice lost >20% initial BW (Fig. 4B right panels). The microneutralization assay indicated that 7A3 exhibited higher IC_50_ against B.1.617.2 than B.1.351, while 8A2 completely lost neutralization against B.1.617.2 (Fig. 3B). Consistent with the microneutralization results, it required 7A3 at a dose of 10 mg/kg to protect 75% of K18-hACE2 mice from the lethality of B.1.617.2; whereas mice receiving the same dose of 8A2 all succumbed to death (Fig. 4B right panels). The K18-hACE2 mice that survived the lethal B.1.617.2 infection developed comparable spike-specific IgG titers in the 7A3 group (IgG GMT of 5120) and the 2-in-1 cocktail group (IgG GMT of 3620) at 4 weeks later (Fig. 4C).

**Fig. 4.**
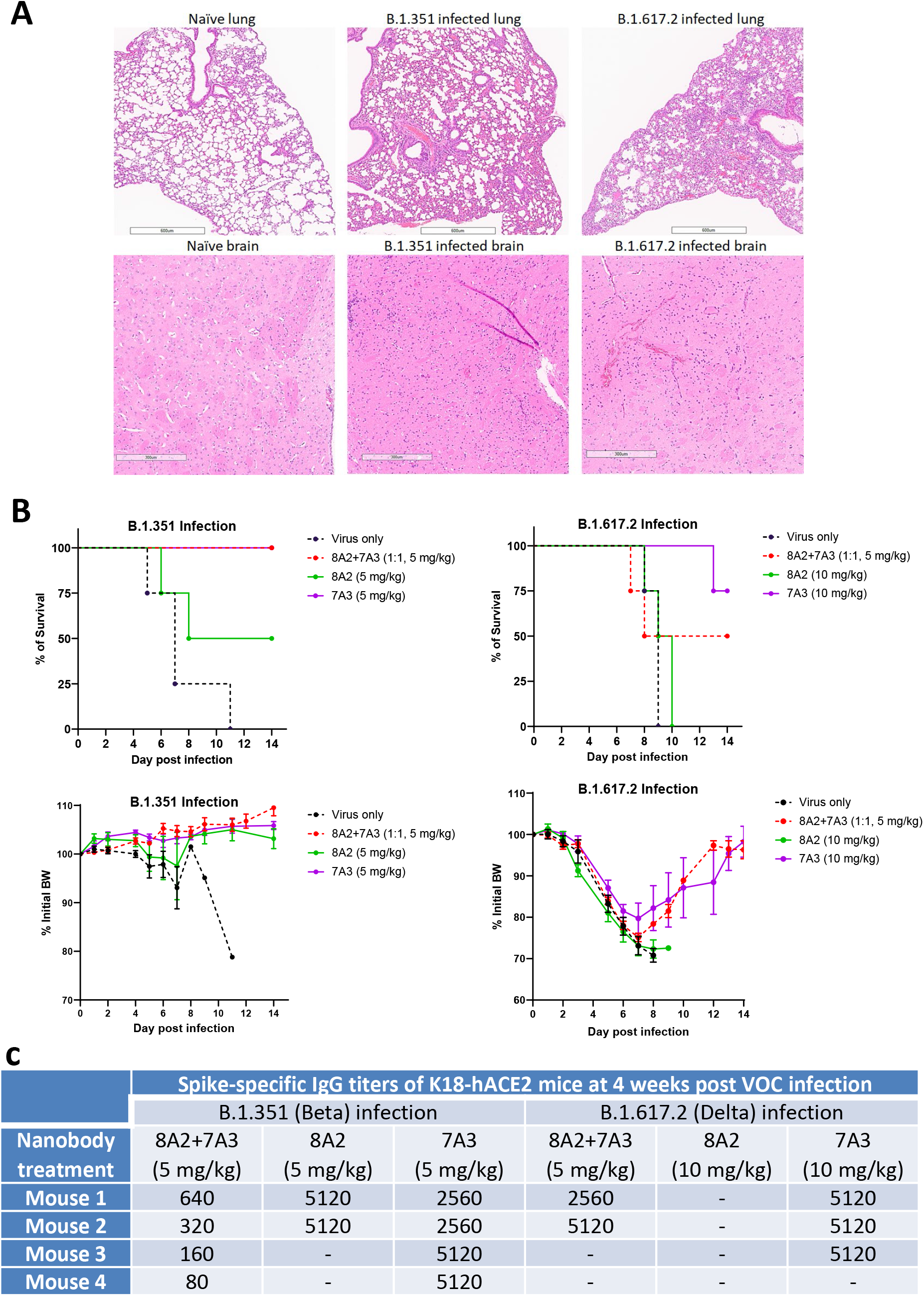
Protection of the K18-hACE2 mice infected with a lethal dose of B.1.351 or B.1.617.2 variant. (**A**) The histology of lung and brain tissues harvested from K18-hACE2 transgenic mice on day 6-7 following infection of live SARS-CoV-2 B.1.351 or B.1.617.2 variant virus. **(B)** K18-hACE2 mice (n=4/group) were injected intraperitoneally with nanobody 7A3, 8A2 or 2-in-1 cocktail at indicated doses followed by a lethal infection of B.1.351 or B.1.617.2 strain. Mortality and body weight (BW) were monitored for 2-week post infection. **(C)** Spike-specific IgG titers of K18-hACE2 mice surviving the lethal B.1.351 or B.1.617.2 infection. -: mice died during infection.

### Structure complexes of the SARS-CoV-2 spike and nanobodies

To precisely characterize the binding of nanobodies on SARS-CoV-2 spike protein, we solved the cryo-EM structures of the hexapro ectodomain in complex with 8A2 V_H_H and with 7A3 V_H_H and of the ternary complex, including both V_H_H nanobodies (Table S4). The overall resolution estimates for the complexes were 3.4, 3.8, and 2.4 Å, respectively. The higher resolution for the ternary complex can be attributed to the instrument used to collect data, the larger dataset, but also to reduced movement of the RBD locked by two nanobodies as evidenced by the local resolution map (Fig. S7). Local refinement of the regions of interest on the ternary complex involving the nanobodies and the RBD yielded three partial maps focused on these areas at overall resolutions of 3.4, 2.7, and 2.6 Å, respectively (Figs. S7-9). The final complex structure showed 8A2 V_H_H indeed interacts with the epitope directly overlapping the site for ACE2 binding when the RBD is in the up mode (Fig. 5A and Fig. S7). Interestingly, 8A2 has the most extended CDR3 loop (21 aa) that penetrates the N-terminal part of the RBD for direct binding of ACE2. 8A2-RBD interaction shows minor variations in the two poses observed (8A2-RBD_B and 8A2-RBD_C) with a common pattern involving sections from three β strands forming the nanobody’s framework extending from Thr33-Ala40; Gly44-Trp47; Tyr95-Thr101 and Gln120. 7A3 V_H_H binds an area that is distinct from the 8A2 site and as demonstrated in RBD_B and RBD_C. 7A3 V_H_H can bind the RBD in both up (RBD_B and RBD_C) and down (RBD_A) modes (Figs. S7 and S9). 7A3-RBD interaction is largely dominated by the interaction of its CDR3 (Tyr102 to Gly113) and an extended cluster of aromatic and hydrophobic residues including Phe37, Leu45, Trp47, Tyr59, Trp101, Tyr107 and Trp112, forming a well structure motif interacting with RBD residues Tyr508, Phe374, Phe377 and Tyr369, and the carbonyl of Phe374 (Fig. 5B and 5C). This motif ends at the tip of the CDR by Trp105. This pattern is common to all three poses of 7A3 found in the structure. Interestingly, nanobody 7A3 can bridge two RBDs in up-down or down-up conformation. However, the up-up disposition does not allow 7A3 bridging. When the RBD is in its down mode (RBD_A), both 7A3-A and a nearby 7A3 V_H_H (7A3-B) can bind the RBD-A while 7A3-B interacts with the region partially overlapping the ACE2 binding site. This interaction does not interfere with 8A2 binding, which is only observed in the up conformation. This observation is consistent with our biophysical cross-competition data on Octet, showing 7A3 can also inhibit ACE2 binding to the RBD. Furthermore, 7A3_A forms interactions with spike chains B and C. Ser17 and Thr69 engage Asp28C and Gly413C carbonyl, while Asn84 engages Asp427C and Arg19 forms a salt bridge with Tyr380C effectively forming a bridge between the two RBDs A and C. Even more remarkable is the interaction network formed at the tip of the 7A3_A nanobody, deep buried in the structure of the spike, where 7A3’s Trp105 and Gln104 engage Asp985/B, Glu988/B, Pro987B and Pro384/A in the S2 subunit of the spike. L452, a key interacting residue with the 8A2 V_H_H (Fig. Fig. 5B and 5C), is critical for the nanobody binding because the mutation of this residue in the B.1.617.2 spike might contribute to the loss of the 8A2 binding (Fig. 1D), E484, another key residue that is mutated in the B.1.617.2 spike, might have less impact on 8A2 binding. Neither L452 nor E484 is involved in the 7A3 V_H_H binding; therefore, 7A3 has broad cross-reactivity with all the major variants. Furthermore, both nanobodies insert into the relatively glycan-free regions of the RBD (Fig. 5A upper right panel). Together, our structure analysis based on cryo-EM maps supports that 8A2 and 7A3 bind two distinct sites on RBD, whereas 8A2 directly disrupts the ACE2 binding on the viral receptor recognition motif and 7A3 binds a conserved site and a buried site on the spike regardless of the conformational state of the RBD.

**Fig. 5.**
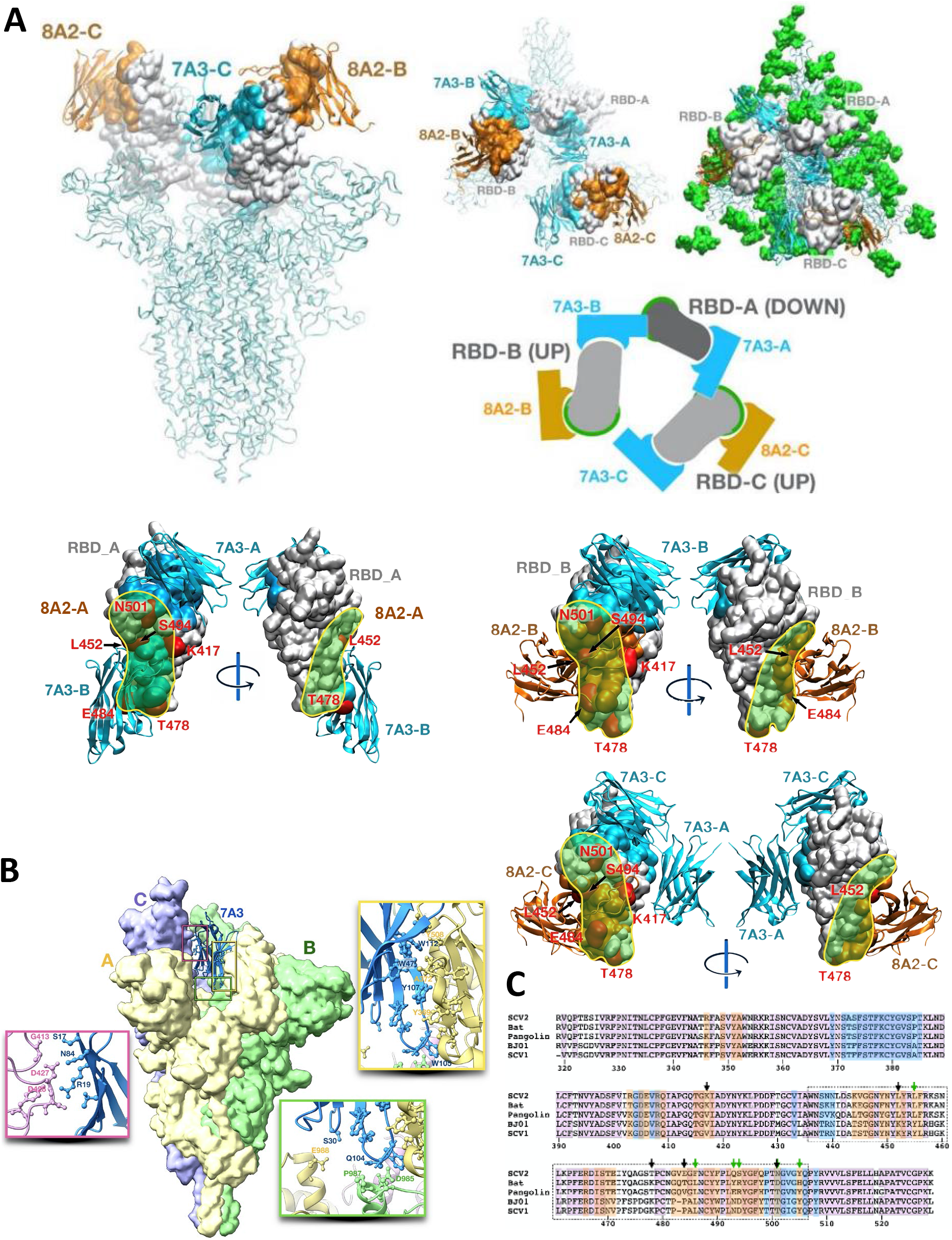
Cryo-EM structure of 7A3 and 8A2 nanobodies with SARS-Cov-2 spike. (**A**) The structure models based on the EM images show that 8A2 and 7A3 bind two distinct sites on the RBD. 8A2 directly interferes with the ACE3 binding to the RBD. 7A3 binds a distinct conserved site on the RBD regardless of its conformational mode. 7A3 partially overlaps the ACE2 binding loci when bridging two RBDs. This interaction does not interfere with 8A2 binding, which is only observed in the up conformation. The mutations on the RBD derived from alpha (E484K, S494P, N501Y), beta (K417N, E484K, N501Y), gamma (K417T, E484K, N501Y), and delta (K417N, L452R, and T478K) were indicated. Green spheres, glycans. (**B**) The unique 7A3 binding pattern. The tip of 7A3 (7A3_A) is deeply buried in the structure of the spike, where the V_H_H nanobody engages Asp985/B, Glu988/B, Pro987B and Pro384/A in the S2 subunit of the spike protein outside the RBD region. (**C**) Nanobodies epitopes coverage as determined from the experimental structure using a 5A contact cutoff. Sequences of coronavirus RBD for Bat_RaTG13 (A0A6B9WHD3); Human BJ01 (Q6GYR1); Pangolin (A0A6G6A2Q2); SARS-CoV-2 (P0DTC2) and SARS-CoV-1 (P59594) are presented for comparison purpose. Degree of sequence identity is indicated by background pink hue. 7A3 contact is indicated in blue and 8A2 in orange. Arrows indicate mutation sites of concern (black) and key ACE2 contact residues (green). RBM is indicated by the dotted box. Sequence alignment and colorization was performed using Jalview version 2.11.1. Alignments were performed using ClustallO and colorization by Blosum62.

## Discussion

In the present study, we constructed large V_H_H phage libraries from dromedary camels and isolated potent neutralizing nanobodies against SARS-CoV-2. Epitope mapping and cryo-EM reveals two distinct epitopes on the RBD for neutralization of SARS-CoV-2. The one recognized by 7A3 V_H_H is conserved between SARS-CoV-2, SARS-CoV and other close coronaviruses (Fig. 5B), and the other recognized by 8A2 V_H_H is unique for SARS-CoV-2. The 8A2 V_H_H directly interferes with the ACE2 binding on the spike protein when the RBD is in its up mode, whereas the 7A3 nanobody binds the RBD in both up and down modes. Consequently, cocktail treatments of nanobodies 7A3 and 8A2 have potent neutralization activities against multiple SARS-CoV-2 variants. Interestingly, 7A3 that binds a conserved RBD epitope and a buried site in the spike can protect mice from B.1.617.2 variant infection. The key contribution and finding of this study are our creation of large dromedary camel V_H_H phage libraries and using the novel camel libraries for isolation of high-affinity V_H_Hs such as 7A3 and 8A2 targeting SARS-CoV-2 spike, which can neutralize the variants of concern in mice. 7A3 has a unique binding site which is highly conserved and deeply buried in the spike that is even extended to the S2 subunit (P987 and D985) outside the RBD region. Consequently, it has broad neutralization activity against all the major emerging variants, indicating that 7A3 has promising therapeutic potential for treating COVID-19. The unique 7A3 eptiope might be used as a template for designing universal vaccines for current and future variants of SARS-CoV-2 or related coronviruses.

In the K18-hACE2 mouse model, B.1.351 has been shown to be 100 times more lethal than the original Wuhan-Hu-1 strain (*21*). After the treatment with the 2-in-1 cocktail 8A2/7A3 nanobodies, K18-hACE2 mice exhibited no sickness and no body weight loss following the lethal challenge of B.1.351. In addition, 7A3 alone can also protect all the mice from severe syndromes though being less efficient in blocking virus infection than the 2-in-1 cocktail. Interestingly, all the surviving mice can develop anti-viral spike humoral response, which may provide a long-term protection. Furthermore, 7A3 can also protect the mice with B.1.617.2 infection. Our data demonstrate that the neutralizing nanobodies can mitigate the COVID-19 syndrome and prevent death without causing noticeable side effects in mice.

Structure analysis based on cryo-EM shows that 8A2 and 7A3 bind two distinct sites on the RBD. 8A2 directly interferes with ACE2 binding by binding to the RBD in the up conformation. In particular, the long CDR3 loop of 8A2 V_H_H might play an important role in its inhibitory function. However, 7A3 can bind the RBD regardless of its up or down mode. We found that 7A3 V_H_Hs could contribute to the stabilization of the two up arms by bridging the down and up conformations. The unique 7A3 binding pattern might be important for the nanobody to achieve its maximum inhibitory effect on ACE2. We find that the tip of 7A3 (7A3-A) is deeply buried in the structure of the spike outside the RBD region, where the V_H_H nanobody engages Asp985/B, Glu988/B, Pro987B and Pro384/A in the S2 subunit of our cryo-EM complex structure (Fig. 5B). This region is key for the function of the spike, as discussed in the recent work on the spike elongation requiring a stable helical bundle at the core of the spike (*22*). The Asp985-Glu988 seems highly conserved as well, indicative of a preserve function and lack of immune surveillance due to the buried disposition of the region. This arrangement suggests a novel mechanism for the stabilization of the spike by 7A3, by forming four different simultaneous contacts: with RBD_A through its epitope, with RBD_A at the spike core region, with RBD_C through its frame and with the spike chain B at the spike core. This peculiar arrangement requires a spike disposition with one RBD-down which may inhibit the full engagement of the spike and serve as a kinetic hindrance. The combination with 8A2, or other motifs blocking the RBM region of the chains in the Up disposition may provide, together, a novel mechanism to maximize efficacy of the intervention by addressing both the stability of the spike, potentially affecting the kinetics of the spike rearrangement while targeting a low mutability spot in the spike and blocking ACE2 binding.

In conclusion, this study characterized two novel nanobodies, 7A3 broadly and 8A2 at a lesser extent neutralizes SARS-CoV-2 and variants. 7A3 binds a highly conserved epitope on the RBD and can neutralize all the major variants that have been emerging so far, including B.1.617.2. Based on these findings, we believe that our leading nanobodies have great therapeutic potential as passive immunotherapy as well as valuable building blocks for developing multi-domain (*23*) and multi-specific drugs for the challenge posed by current and future emerging variants. The characterization of the 7A3 epitope sequence and structure might provide new insights for vaccine design broadly targeting SARS-CoV-2 variants as well as similar coronaviruses in the future.

## Acknowledgments

We thank Nicole Doria-Rose and Kizzmekia Corbett at the Vaccine Research Center, National Institute of Allergy and Infectious Diseases (Bethesda, Maryland), Alex Compton, Alan Rein and Kin Kui Lai at the National Cancer Institute (Frederick, Maryland) for providing the SARS-CoV-2 pseudovirus and the protocols used in the present study, Zina Itkin and Paul Shinn for compound management support for CPE assays, Hui Guo for CPE assay data analysis, and Wei Zheng for scientific guidance and discussions at the National Center for Advancing Translational Sciences (Rockville, Maryland), Yihong Ye and Qi Zhang at National Diabetes and Digestive and Kidney Diseases (Bethesda, Maryland) for helpful discussion in SARS-CoV-2 cellular assays and animal testing. The content of this publication does not necessarily reflect the views or policies of the Department of Health and Human Services, nor does mention of trade names, commercial products, or organizations imply endorsement by the U.S. Government.

The V_H_H nanobodies targeting SARS-CoV-2 spike protein are the subject of the pending patent application no. PCT/US2021/056548 and are available for license in certain fields of use to qualified candidates. Please contact the leading corresponding author Dr. Mitchell Ho at if you are interested in pursuing a license.

## Funding

NIH Intramural Targeted Anti-COVID-19 (ITAC) Program (ZIA BC 011943 to M.H. and ZIA ES 103341 to M.J.B.)

Intramural Research Program of NIH, Center for Cancer Research (CCR) National Cancer Institute (NCI) (Z01 BC010891 and ZIA BC010891to M.H.)

NCI CCR Antibody Engineering Program (ZIC BC 011891 to M.H.) NIEHS Cryo-EM Core (ZICES10326 to M.J.B.)

Intramural Research Program of the National Center for Advancing Translational Sciences, NIH (to C.Z.C. and M.X.)

FDA/CBER intramural SARS-CoV-2 fund (to H.X.).

National Cancer Institute, National Institutes of Health, Department of Health and Human Services, Contract No. 75N91019D00024 (to R.C. and D.E.)

## Author contributions

Conceptualization: M.J.B., H.X., and M.H.

Methodology: J.H., H.J.K., R.C., C.Z.C., K.J.B., Z.D., D.L., H.R., J.Z., V.P.D., N.M., M.J.B., D.E., U.O.R., M.X., M.J.B., H.X., and M.H.

Investigation: J.H., H.J.K., R.C., C.Z.C., K.J.B., Z.D., D.L., T.L., J.Z., V.P.D., N.M., M.J.B., U.O.R., H.X., and M.H.

Visualization: J.H., R.C., M.J.B., H.X., and M.H. Funding acquisition: M.J.B., H.X., and M.H.

Project administration: J.H., H.J.K., R.C., N.M., D.E., M.X., M.J.B., H.X., and M.H. Supervision: M.J.B., H.X., and M.H.

Writing – original draft: J.H., R.C., M.J.B., H.X., and M.H.

Writing – review & editing: J.H., R.C., C.Z.C., M.J.B., H.X., and M.H.

## Data and materials availability

All data are available in the main text or the supplementary materials.

