## Supplementary materials for "Camel nanobodies broadly neutralize SARS-CoV-2 variants"

**This PDF file includes:**

Materials and Methods
Figs. S1 to S9
Tables S1 to S4

### Materials and Methods

#### Regents related to COVID-19

The following reagents were obtained through BEI Resources, NIAID, NIH: SARS-Related Coronavirus 2: (1) Isolate USA-WA1/2020 bearing 614D, NR-52281 (deposited by the Centers for Disease Control and Prevention); (2) Isolate New York-PV09158/2020 bearing 614G, NR-53516; (3) Isolate USA/CA_CDC_5574/2020 (B.1.1.7 or Alpha), NR-54011 (deposited by the Centers for Disease Control and Prevention); (4) Isolate hCoV-19/South Africa/KRISP-EC-K005321/2020 (B.1.351 or Beta), NR-54008 (contributed by Alex Sigal and Tulio de Oliveira); (5) Isolate hCoV-19/Japan/TY7-503/2021 (Brazil P.1 or Gamma), NR-54982 ; (6) Spike Glycoprotein (Stabilized) from SARS-Related Coronavirus 2, Wuhan-Hu-1 with C-Terminal Histidine Tag, Recombinant from Baculovirus, NR-52396. (7) Vector pCAGGS containing the SARS-Related Coronavirus 2, Wuhan-Hu-1 (or USA-WA-1) Spike Glycoprotein Receptor Binding Domain (RBD), NR-52309. (8) Isolate B.1.617.2 (Delta) Lot 3002648422 was kindly provided by Dr. Bin Zhou (the Centers for Disease Control and Prevention). All live viruses were amplified in Vero E6 (ATCC, CRL-1586) or TMPRSS2-E6 (BPS Bioscience, #78081) and aliquots were stored at -80°C until use.

#### Construction of camel nanobody libraries

The construction of the camel library followed our previous protocol (*23*) for a shark $V_{NAR}$ phage library with some modifications. The PCR was performed with the forward primer CALL001 (5-GTCCTGGCTGCTCTTCTACAAGG-3) and reverse primer CALL002 (5-GGTACGTGCTGTTGAACTGTTCC-3). The amplified $V_HH$ fragments were cloned into the phagemid vector backbone pComb3X to generate six individual $V_HH$ phage libraries with the size of $10^{10}$ for each library. The total diversity of six $V_HH$ libraries used in the present study is above $10^{11}$.

**Phage panning and ELISA**

Phage panning generally follows our published protocols (*24, 25*). In one approach, Maxisorp immunotubes (Thermo Scientific) were coated with SARS-CoV-2 RBD (Sino Biological, 40592-V08B1) for the first and second round of panning and SARS-CoV-2 S trimer (BEI, NR52396) for the 3rd and 4th rounds of panning. In another approach, SARS-CoV-2 RBD was used for the first, second, and third round of panning and SARS-CoV-2 S trimer for the 4th round.

After phage panning, single colonies were picked for monoclonal ELISA. Maxisorp 96-well plate (Fisher Scientific, 12565136) was coated with SARS-CoV-2 RBD, S trimer, or SARS-CoV-1. Phage ELISA generally follows previous protocols (*24, 25*), and results were read using a spectrophotometer (Molecular Devices) at 450 nm.

**Analysis of complementarity-determining regions (CDRs).**

Sequence alignment of SARS-CoV-2 neutralizing $V_HHs$ using Clustal Omega Program with IMGT in bold, Kabat italicized, and Paratome underlined. IMGT and Kabat were determined using IgG Blast (https://www.ncbi.nlm.nih.gov/igblast/), and Paratome was determined using Ofran Lab service (http://ofranservices.biu.ac.il/site/services/paratome/).

**Affinity measurement and competition assay by Octet**

The binding kinetics and competition assay were determined using the Octet RED96 system (FortéBio) at the Biophysics core at NHLBI, NIH (Bethesda, MD). For binding kinetics, the SARS-CoV-2 original or mutant variant was immobilized onto Ni-NTA sensor tips (Fortebio).

The antigen-coated tips were then dipped into PBS to stabilize the curve, transferred into 25 nM $V_H$H-hFc for association, and finally dipped into PBS for dissociation. Raw data was processed using Octet Data Analysis Software 9.0 to determine a/the $K_D$ value.

For the competition assay, the SARS-CoV-2 RBD-His was immobilized onto Ni-NTA sensor tips. The resulting RBD-coated tips were then dipped into either PBS or 500 nM first nanobodies. After loading, the sensor tips were briefly incubated in PBS before being dipped into wells containing 500 nM of the competing nanobody, followed by a dissociation step in PBS. Raw data was processed using Octet Data Analysis Software 9.0. Residual binding was calculated as follows: (response signal from the second ligand in the presence of first ligand/response signal from the second ligand in the absence of the first ligand) x 100.

**ACE2 inhibition assay**

A Ni-NTA ELISA plate (brand) was coated with ACE2-His. $V_H$H-hFc was incubated with varying concentrations of RBD-mFc starting from 1 μg/ml with 1:3 dilutions. The $V_H$H-hFc and RBD-mFc mixture was then added to the ACE2-His coated plate, and binding was detected using a goat anti-mouse Fc HRP conjugate (Jackson ImmunoResearch). Results were read at the wavelength of 450 nm using a spectrophotometer (Molecular Devices).

**Flow cytometry**

The spike coding sequences for SARS-CoV Urbani and SARS-CoV-2 in pcDNA3.1 (+) plasmid were provided by Alex Compton (NCI, Frederick). These sequences were then codon-optimized for human cell expression, followed by a 5′ Kozak expression sequence (GCCACC), 3′ tetra-glycine linker, and a FLAG-tag (DYKDDDDK)(*26*). A431, a human epidermoid carcinoma cell line, was transfected with either pcDNA3.1 (+)-SARS-CoV-spike vector or pcDNA3.1 (+)-

SARS-CoV2-spike vector by either FuGENE HD (Promega) or Lipofectamine 3000 (ThermoFisher). The A431 clones with high viral spike protein expression on the cell surface were sorted by flow cytometry using the CR3022 antibody for SARS-CoV and the 2019-nCoV antibody (SinoBiological) for SARS-CoV-2. Either A431-CoV-2-S or A431-CoV-S cell line was stained with camel $V_H$H-hFc followed by goat anti-human IgG-APC (Jackson ImmunoResearch). CR3022 antibody was used as the positive control. Data was collected using BD FACSCanto$^{TM}$ II Cell Analyzer.

**Pseudoviral neutralization assays**

Three pseudo viral neutralization assays were conducted independently in different laboratories to validate the anti-viral activities of the camel $V_H$H nanobodies. In the first assay, called lentivirus-based pseudovirus infection assay, HEK293T cells expressing human ACE2 (HEK293T-ACE2) (provided by Nicole Doria-Rose and Kizzmekia S. Corbett, Vaccine Research Center, NIAID) were seeded in 96-well plates. The appropriate volume of SARS-CoV-2 or SARS-CoV spike pseudovirus supernatant was used to produce a luciferase signal 1000x higher than the baseline. $V_H$H-hFc fusion proteins were prepared in 12 point 2-fold serial dilutions starting with 50 μg/ml for SARS-CoV-2 and 100 μg/ml for SARS-CoV. For the "2-in-1 cocktail", $V_H$H-hFcs were prepared in 12 point 2-fold serial dilutions starting with 25 μg/ml each nanobody for a total of 50 μg/ml. Nanobodies and viruses were mixed for 45 mins before adding to HEK 293T-hACE2 cells. After incubation for 72 hrs, the luciferase signal was measured by the plate reader. All experiments were performed in triplicate. The infectivity was calculated from the luciferase activity of different groups normalized by cells with virus only group (100%). The graphs were made, and the $IC_{50}$s and $IC_{90}$s were calculated by GraphPad Prism.

In the second assay called pseudotyped particle (PP) entry assay, SARS-CoV-2-S Wuhan-Hu-1, B.1.351, P.1, and B.1.1.7 PPs were purchased from Codex Biosolutions (Gaithersburg, MD) and were produced using a murine leukemia virus (MLV) pseudotyping system (*27*). The variant spike sequences clones were: B.1.351 (L18F, D80A, D215G, del242-244, K417N, E484K, N501Y, D614G, and A701V), B1.1.7 (del69-70, del144, N501Y, A570D, D614G, P681H, T716I, S982A, and D1118H), and P.1 (L18F, T20N, P26S, D138Y, R190S, K417T, E484K, N501Y, D614G, H655Y, T1027I, V1176F). Expi293F cells with stable expression of human ACE2 (HEK293-ACE2) were generated at Codex BioSolutions (Gaithersburg, MD). For the PP entry assay, HEK293-ACE2 cells were seeded in 384-well microplates (Greiner BioOne). The cells were incubated at 37 °C with 5% $CO_2$ overnight. $V_H$H-hFc fusion proteins at 120 µg/ml were titrated 1:3 in DPBS for 12 concentration points and were added to cells in triplicates. Cells were incubated with nanobodies for 1 hr at 37 °C 5% CO2 before SARS-CoV-2-S PP was added. The plates were then spinoculated by centrifugation at 1500 rpm (453 x g) for 45 mins and incubated at 37 °C for 48 hrs to allow cell entry of PP and expression of the luciferase reporter. After the incubation, the supernatant was removed with gentle centrifugation using a Blue Washer (BlueCat Bio). Then Bright-Glo Luciferase detection reagent (Promega) was added to assay plates and incubated for 5 mins at room temperature. The luminescence signal was measured using a PHERAStar plate reader (BMG Labtech). Data was normalized with wells containing PPs as 100% and wells containing no PP (media control) as 0%. All nanobodies were also assessed for cytotoxicity as a counter assay using the same cell treatment and incubation protocol, omitting the PP addition, and assaying for ATP content with ATPLite (PerkinElmer) cytotoxicity kit.

The third assay is called the pseudovirus fluorescence reporter assay (*28*).

**Live SARS-CoV-2 neutralization assay**

Two live virus assays were conducted independently in different laboratories. The first assay is called SARS-CoV-2 cytopathic effect (CPE) assay. The CPE assay was performed at the Southern Research Institute (Birmingham, AL) (*29*). Briefly, $V_H$H-hFc fusion proteins were titrated in PBS and acoustically dispensed into 384-well assay plates at 600 nL/well. Next, cell culture media (MEM, 1% Pen/Strep/GlutaMax, 1% HEPES, 2% HI FBS) was dispensed at 5 µL/well into assay plates and incubated at room temperature. Vero E6 previously selected for high ACE2 expression was dispensed to the plate at 4000 cells/well in 10 µl media. The cells were incubated with nanobodies for 30 min, before SARS CoV-2 (USA_WA1/2020) was inoculated at 0.002 M.O.I. in 15 µl/well media. Assay plates were incubated for 72 hr at 37 °C, 5% $CO_2$, 90% humidity. Then, CellTiter-Glo (Promega) was dispensed, incubated for 10 mins at room temperature, and luminescence signal was read on an EnVision plate reader (PerkinElmer). Data was normalized with wells containing virus as 0% CPE rescue and wells without virus (media control) as 100% CPE rescue.

The second assay is a live SARS-CoV-2-based microneutralization assay. Virus titers were determined using an ELISA-based 50% tissue culture infectious dose ($TCID_{50}$) method (*30*). Vero E6 cells were pre-seeded in 96-well tissue culture plates overnight. Next day individual $V_H$H-hFc fusion proteins or 2-in-1 mixtures were serially diluted and were incubated with $10^2$ $TCID_{50}$ of the live virus at room temperature for 1 hr. The virus-nanobody mixtures were then added to Vero E6 cells pre-seeded in 96-well flat-bottom tissue culture plates and incubated at 37°C, 5% $CO_2$ for another 48 hr. The residual virus was detected using in-house developed SARS-CoV-2 specific rabbit polyclonal antibodies (*30*) followed by goat anti-rabbit IgG with horseradish peroxidase conjugate (Invitrogen). All $V_H$H-hFc fusion proteins were tested at the starting concentration of 120 nM or 60 nM in the 2-in-1 mixtures.

**Animal testing**

All procedures were performed according to the animal study protocols approved by the FDA White Oak Animal Program Animal Care and Use Committee. Hemizygous 2B6.Cg-Tg(K18-ACE2)2Prlmn/J (K18-hACE2) transgenic mouse strain (JAX) was bred in the FDA White Oak vivarium and were genotype confirmed before the experiments. All subsequent live virus infection experiments were conducted in the FDA animal biosafety level (ABSL) 3 laboratory.

K18-hACE2 mice infected with the hCoV-19/South Africa/KRISP-EC-K005321/2020 (B.1.351, Beta) or B.1.617.2 (Delta)variant were humanely euthanized after they reached the moribundity on day 6-7 post infection. Lung and brain tissues were harvested and fixed in 10% neutral buffered paraformaldehyde for at least two weeks before histology. Fixed lung and brain tissues were embedded in paraffin and were sectioned followed by the standard hematoxylin and eosin (H&E) staining (Histoserv). Uninfected mouse lung and brain were used as negative control. H&E images were captured using Leica Aperio AT2 slide scanner (Histoserv) and viewed using Aperio ImageScope DX clinical viewing software (version 12.4.3.5008). Adult K18-hACE2 mice were injected with individual $V_H$H-hFc fusion proteins (7A3 and 8A2) at 5 or 10 mg/kg or the 2-in-1 mixture at 5 mg/kg via the intraperitoneal route. Approximately 2 hrs later, mice were anesthetized under isoflurane and were intranasally inoculated with B.1.351 or B.1.617.2 at $10^2$ $TCID_{50}$ per mouse. Body weight (BW) and mortality were monitored daily for up to 14 days post-infection. All efforts were made to minimize animal suffering, and mice reaching predefined humane endpoints (e.g., 30% BW loss) were immediately euthanized.

The mice that survived the infection were tail bled at approximately four weeks post-infection, and sera were collected for spike-specific IgG ELISA as described before (*30*). All sera were heat-inactivated at 56°C for 30 min before ELISA. Mouse sera were serially diluted in PBS (pH7.4) and were added to recombinant spike pre-coated 96-well microtiter plates (*30*). Bound mouse IgG was probed using peroxidase-conjugated goat anti-mouse IgG (Invitrogen) followed by One-step TMB substrate (ThermoFisher). The endpoint IgG titers were the reciprocals of highest serum dilutions that yielded > two-fold of optical density (OD) than that of PBS blank at 450 nm.

**Cryo-EM specimen preparation**

SARS-CoV-2 S protein ectodomain (spike), hexapro variant, was kindly provided by Dominic Esposito at NCI Frederick (*31*). Complexes of spike with 7A3 and/or 8A2 $V_H$Hs were prepared by mixing the components at a spike trimer to nanobody molar ratio of 1:6. The final concentration of spike trimer was 3 μM in PBS at pH 7 with the addition of 5 mM Imidazole. Because the 8A2 stock was too diluted, complexes involving this nanobody were prepared at 0.5 μM spike trimer followed by a 6-fold concentration using a 10 kDa cutoff centrifugal filter (Amicon Ultra). All complexes were incubated on ice for at least 5 mins prior to grid preparation.

Specimens were plunge frozen on customized support grids consisting of C-flat R 1.2/1.3 (Protochips) supplemented with a 30 nm thick gold layer applied on the grid bar side using a sputterer (Leica ACE-600). Prior to specimen deposition, grids were pretreated on a plasma cleaner (Tergeo, model) in immersion mode with a power of 38 W for a period of 75 s. A 3 μl aliquot of the prepared complex was laid on the surface of the grid inside the chamber of a vitrification robot (Leica EM-GP2) held at 22 °C with an RH of 98%, blotted for 4 s using two layers of filter paper (Whatman Grade 1), immediately plunged into liquid ethane kept at 90 K and transferred to liquid nitrogen for storage.

**Cryo-EM data collection and image processing**

Specimens were imaged on either a TFS/FEI Talos Arctica operated at 200 KeV or a TFS/FEI Titan Krios operated at 300 KeV furnished with a Gatan GIF operated in zero loss mode with a slit of 20 eV. Micrographs were recorded as movies (Supplemental Table 5) on a Gatan K2-Summit or a K3 direct electron detector. Data preprocessing was performed in the context of Scipion 3. Movies were aligned using MotionCorr2 before CTF determination using CTFFIND4.1. Motion corrected dose weighted micrographs were imported into Cryosparc for further processing. Individual molecular images were detected using the Topaz Extract machine learning algorithm. Extracted particles were curated utilizing a combination of 2D classification rounds and Ab-initio Refinement. "Clean" particles were then refined using series of heterogeneous, unsymmetrized homogeneous, C3 symmetrized homogeneous, Global CTF, and Non-Uniform Refinements.

**Molecular Modeling**

8A2 and 7A3 models were obtained using homology modeling tools. The initial model of 8A2 reveals an unusually extended CDR3, so we challenge our model to ensure the quality of our initial guess. YASARA (*32*), I-TASER (*33*), Rosetta (*34*) homology modeling packages were used for this purpose. Local installation of Rosetta (Linux version 2020 08.61146 bundle), I-Tasser (version 5.1), and Yasara (Mac version 20.10.4) were used. Standard scripts and parameters were used for Rosetta and I-Tasser. Yasara homology modeling macro (HM_build), was used allowing 25 templates, ten alternative sequence alignments, and 50 loops per model. Finally, the structures obtained from all the modeling engines were combined using YASARA HM_build macro to form a hybrid model. The hybrid model was allowed to be further refined against all templates using Feedback Restrain Molecular Dynamics (*35, 36*), resulting in a slight improvement of the Z-score from 0.5 to 0.45 for 8A2 and 0.6 to 0.4 for 7A3 but showing a general agreement across the procedures used which confirm the unusual 8A2 CDR3 geometry.

The nanobody models were subjected to molecular dynamics calculations [Gromacs 2021, (*37*)] to build a diverse set of conformations for macromolecular docking [ZDock v 3.0.2 (*38*)] and rigid body fitting to the maps using Chimera version 1.15 (Mac build 42258) (*39*). Multiple orientations were obtained due to the lack of resolution of the initial map. We clustered the poses obtained and performed molecular dynamics simulations starting from a set of hand-curated 240 most promising orientations which were subjected to constrained molecular dynamics simulations, performed using Gromacs-2018-densfit (*40*) on an initial model obtained by fitting PDB ID:6X2B (*38*) to the density using Chimera (*41*), aligning the N-terminal domain structure PDB ID:5X4S (*42*) and RBD structure PDB ID:7EAM which were processed using YASARA homology modeling macro with default parameters to generate missing loops and rigid body aligned to the corresponding sites in the spike reference structure (PDB ID:6X2B) followed by local fitting to density using Chimera. Molecular dynamics trajectories (*37*) were used to generate an epitope map based on the mean contact time of the nanobody with the RBD residues weighted by the correlation coefficient reported by Gromacs-densfit using a 5A cutoff distance. Final structure refinement was performed using PHENIX (1.19.2_4158) (*43*) followed by manual correction using Chimera. A model including glycosylation sites was generated following the PDBID:6x2B glycosylated structure available in the CHARM-GUI repository (http://www.charmm-gui.org/docs/archive/covid19).

```
7A3     QVQLVESGGGSVQPGGSLRLSCVVSGYTSSSRYMGWFRQVPGKGLEWVSGIKRDGTNTYYADSVKGR
8A4     EVQLVESGGGLVQPGESLRLSCEASGFTFSSVYMSWVRQAPGKGLEWISTIHPAGGSTYYADSMKDR
1B5     DVQLVESGGGSVQAGGSLRLSCTGSRYTYSTYCMGWFRQAPGKEEEAVAIINSGGGEPYYGDSVKGRF
8A2     AVQLVDSGGGSVQAGGSLRLSCAASGYTYSICTMGWYRQAPGEGLEWVSGINADGSNTHYTDSVKGR
2F7     QVKLEESGGGSVQSGGSLRLSCTVSRDTNTNINRCMGWFRQAPGKGLETVATINRDGTNTYYTDAVKGR
1H6     AVQLVDSGGGSVQAGGSLNLSCVASGTTLRNGCMAWFRQVPGKEREVVAIIIRATSYTDYADSVKGR

7A3     FTISQDNAKNTVYLQMNSLKPEDTAMYYCAAGSWYNQWGYSMDYWGKGTQVTVSS          122
8A4     FTISRDNAKNTLYLQMNSLKSEDTALYYCIIEALSGYRGPGTQVTVSS                 115
1B5     FTISQDRAKNTVYLQMDGLQPDDTAIYYCVAADSHNSRCYLGRSYVNYWGQGTQVTVSS      126
8A2     FTISRDNAKKTLYLQMNSLKPEDTAIYYCAAHGTYDKYAPCGGFAGTYTYWGQGTQVTVSS    128
2F7     FTISQDNVKNTVYLQMNNLTPEDTGTYICNAMGRGSGSRCDNWDPNYWGQGTQVTVSS       127
1H6     FTISQDNAKNTVYLQMKSLTPEDTATYYCAATLYRVNCAKREFDKWGQGTQVTVSS         123

                IMGT, Kabat, Paratome
```

**Fig. S1.**

Sequence alignment of the top 6 $V_HH$ with CDR regions bolded for IMGT, italicized for Kabat, and underlined for Paratome. The cysteines are highlighted in red. All the $V_HH$ sequences are also listed in provisional patent application no. PCT/US2021/056548.

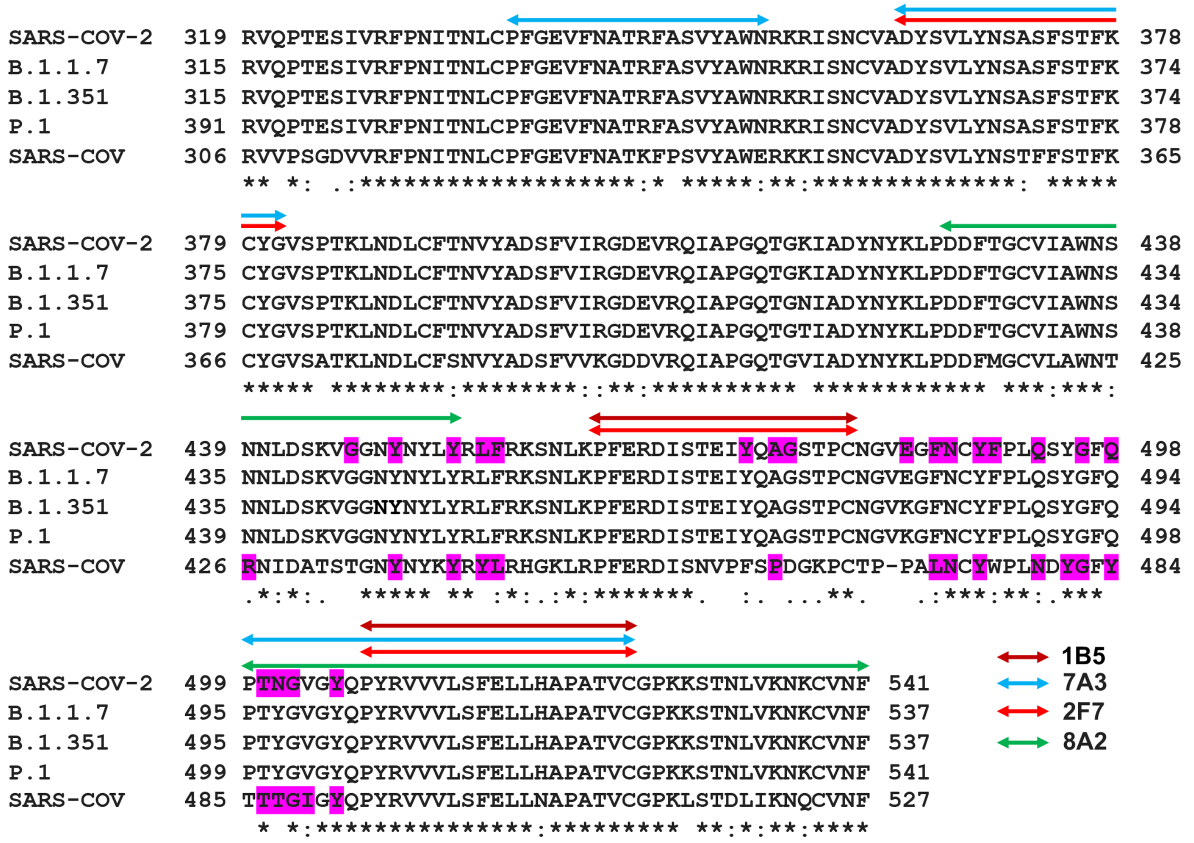


**Fig. S2.**

Epitope mapping of individual anti-SARS-CoV-2 RBD $V_H$Hs. Sequence alignment of RBD region of SARS-CoV-2, SARS-CoV and three identified variant strains of SARS-CoV-2 (B.1.1.7 EPI_ISL_601443, B.1.351 EPI_ISL_700428, and P.1 EPI_ISL_792680). The conserved residues are marked with asterisks (*), the residues with similar properties between variants are marked with the colon symbol (:) and the residues with marginally similar properties are marked with the period symbol (.). The main residues of both SARS-CoV-2 and SARS-CoV RBD region identified previously that interact with ACE2 are shaded in magenta. Arrows of different lengths and colors represent possible positions of contact with RBD.

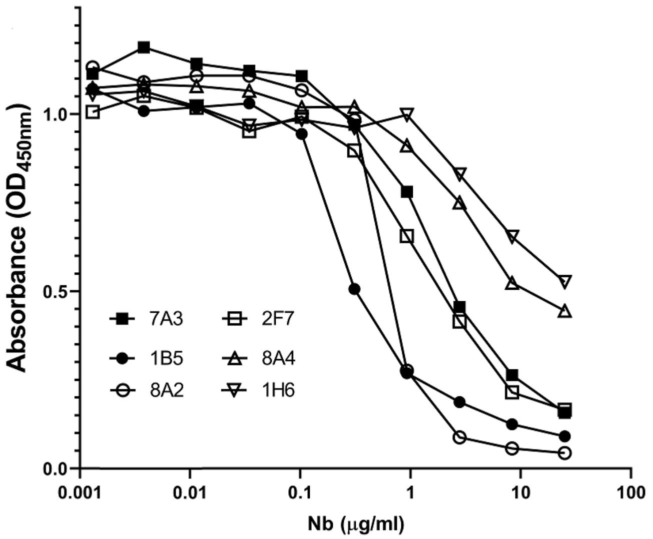


**Fig. S3.**

Inhibition effect of VHH nanobodies against the interaction of the RBD and the human ACE2 protein by ELISA.

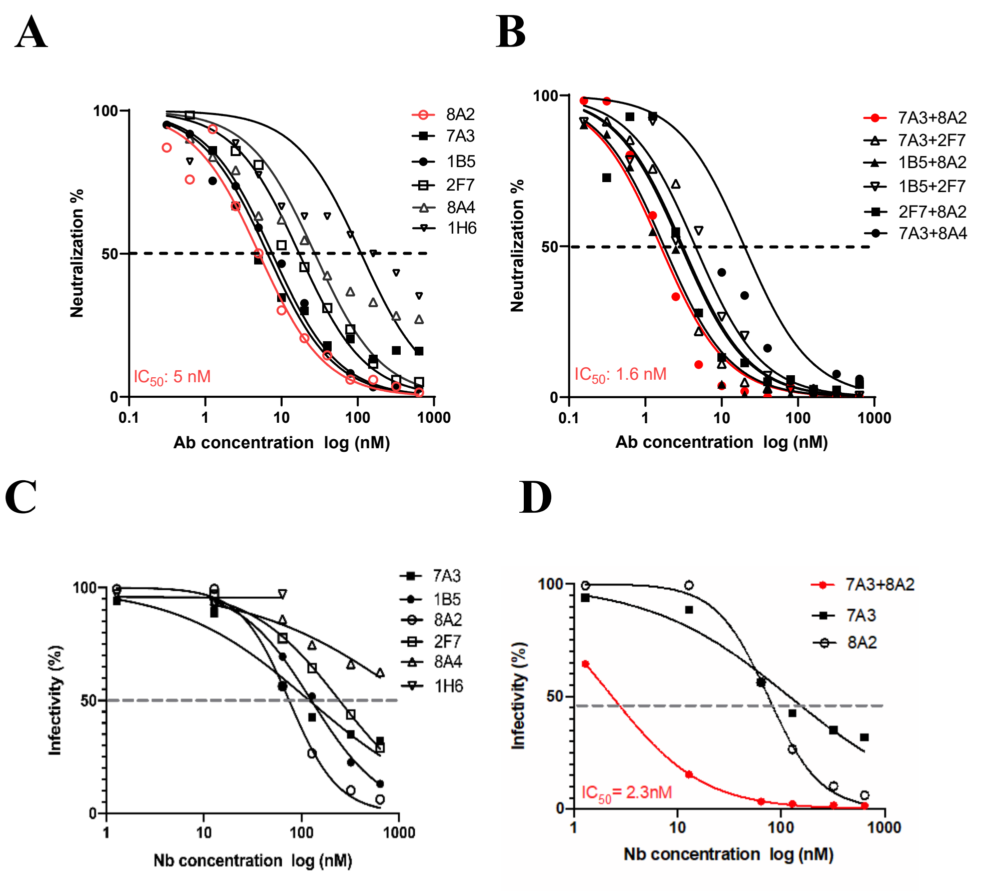


**Fig. S4.**

Neutralization assay on SARS-CoV-2 (Wuhan-Hu-1). 4A-B: Lentivirus-based pseudovirus assay.

4C-D: Fluorescence reporter assay

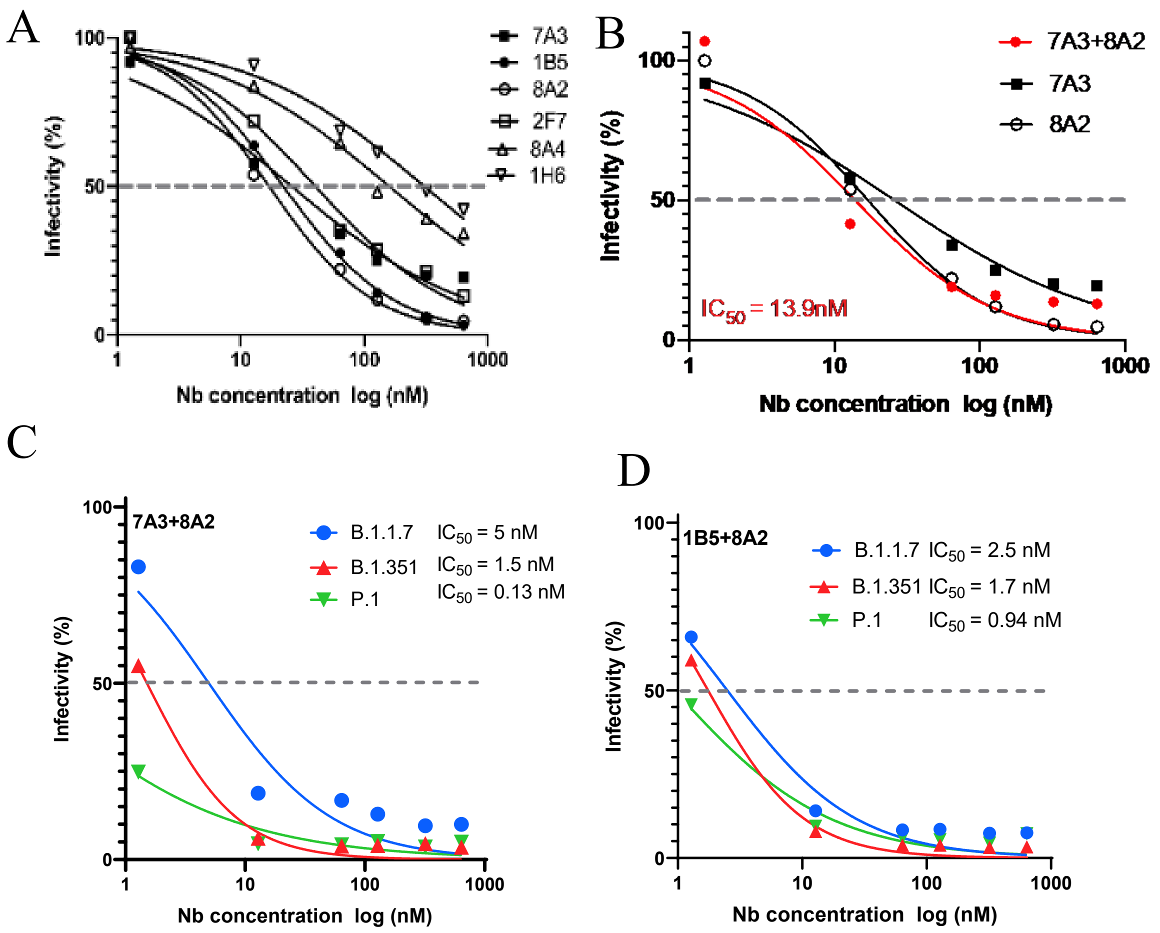


**Figs. S5.**

Neutralization of the mutants of SARS-CoV-2 pseudoviral infection. 5A-B: Fluorescence reporter assays using D614G mutant. 3C-D: Fluorescence report assay using B.1.1.7, B.1.351, and P.1 variants

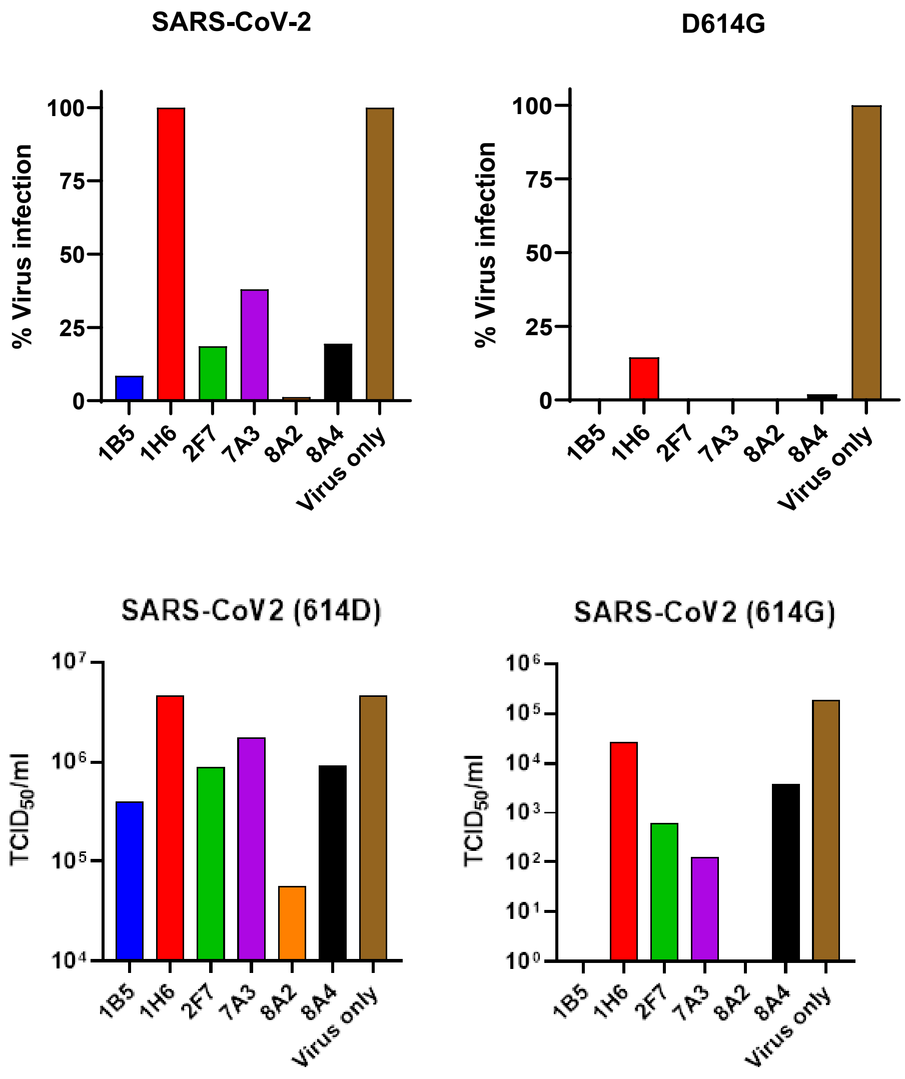


**Fig. S6.**
Prescreening $V_HH$-hFc fusion proteins at 10 µg/ml against live SARS-CoV-2 bearing 614D or 614G. $V_HH$-hFc fusion proteins at the concentration of 10 µg/ml were incubated with $10^2$ $TCID_{50}$ of live SARS-CoV-2 or the 614G variant at room temperature for 1 h. The mixtures were then added to Vero E6 cells pre-seeded in 96-well flat-bottom tissue culture plates and incubated at 37°C, 5% $CO_2$ for 728 h. Residual virus in the supernatants was titrated by $TCID_{50}$ assay.

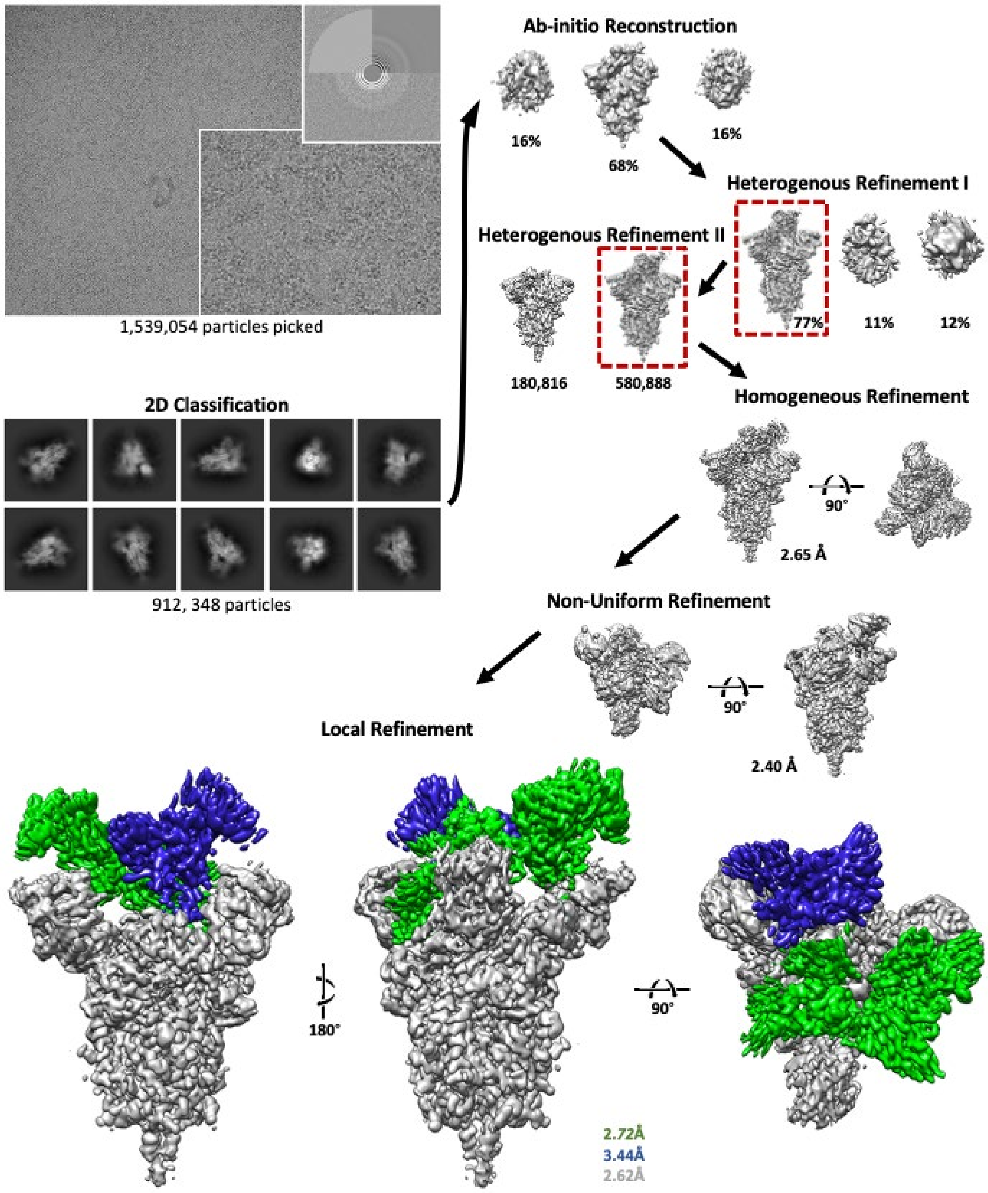


**Fig. S7**.

Cryo-EM image processing workflow for S-protein ectodomain in complex with 7A3 and 8A2. A total of 1,539,054 particles selected from 4,644 micrographs were subject to 2D classification. Class selection yielded 912,348 which were used to generate three initial maps by Ab-initio reconstruction, followed by two cycles of heterogenous refinement and a third non uniform refinement step and a final refinement focused on the RBDs.

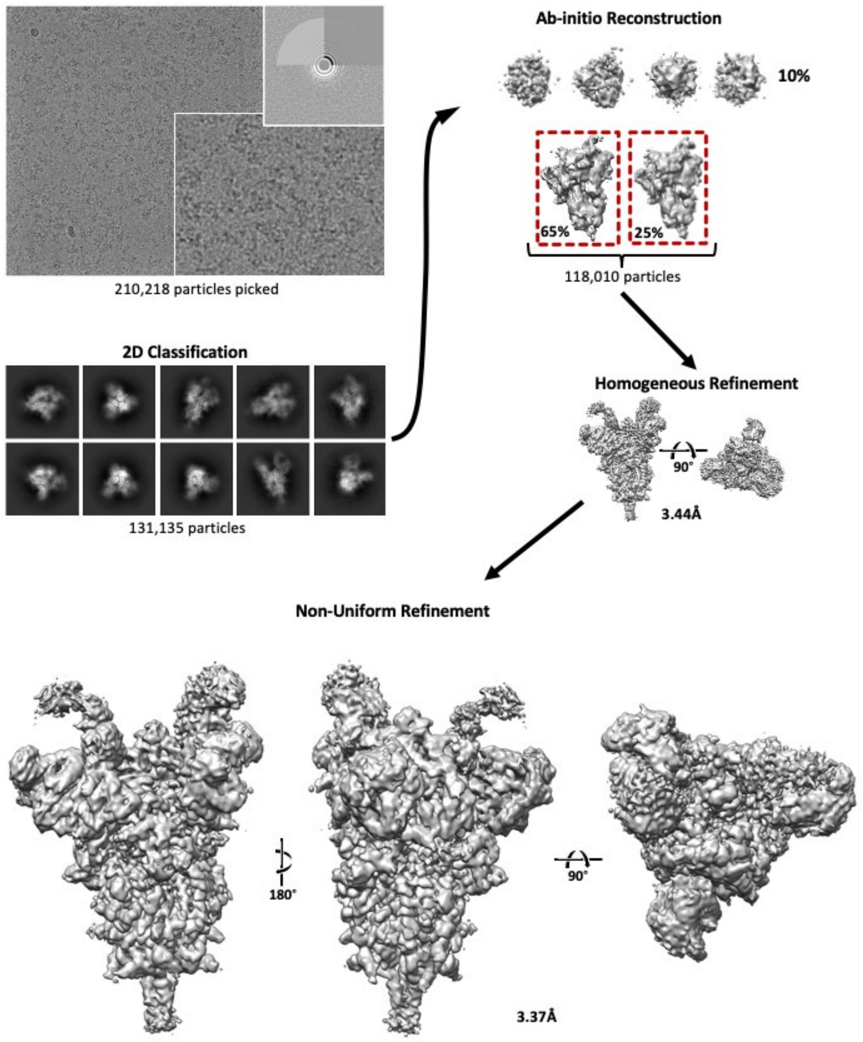


**Fig. S8.**
Cryo-EM image processing workflow for S-protein ectodomain in complex with 8A2. A total of 210,218 particles selected from 2149 micrographs were subject to 2D classification. Class selection yielded 131,135 which were used to generate three initial maps by Ab-initio reconstruction, followed by heterogenous refinement and non-uniform refinement.

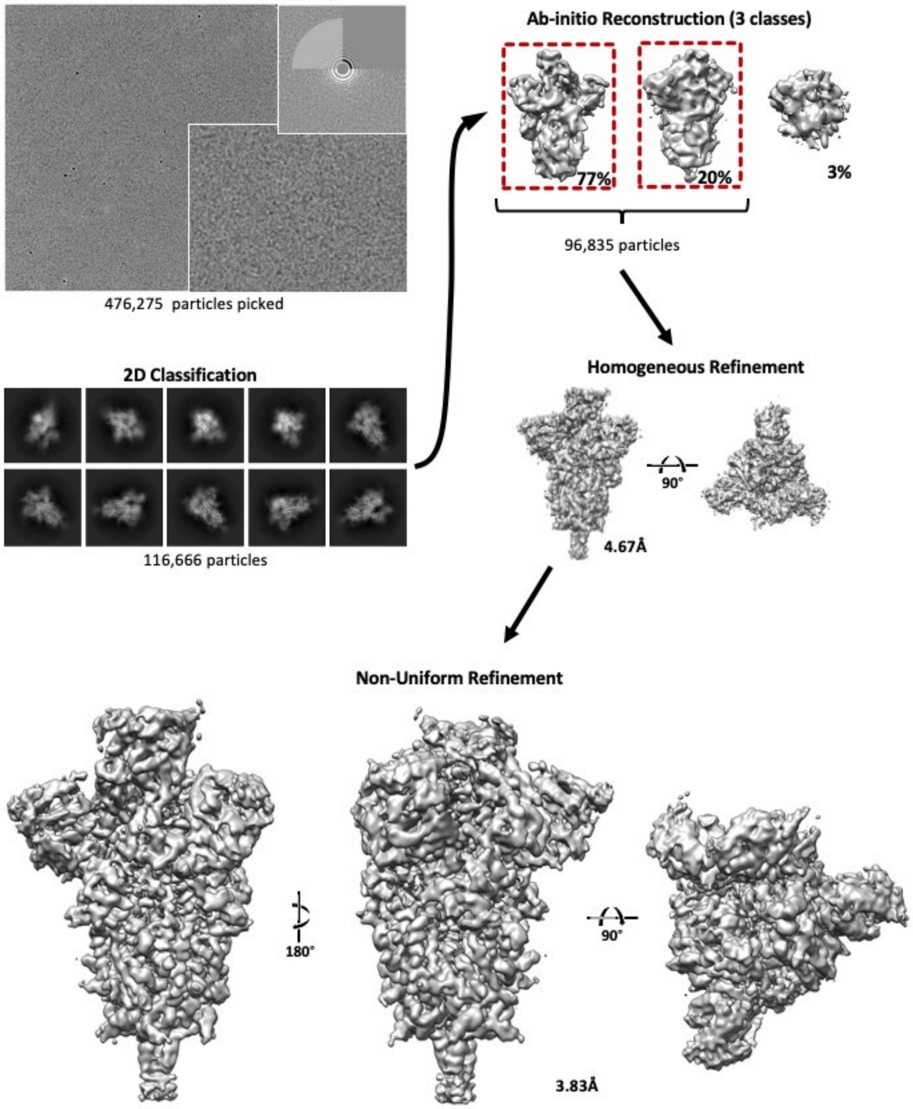


**Fig. S9.**
Cryo-EM image processing workflow for S-protein ectodomain in complex with 7A3. A total of 476275 particles selected from 3198 micrographs were subject to 2D classification. Class selection yielded 116666 which were used to generate three initial maps by Ab-initio reconstruction, followed by heterogenous refinement and non-uniform refinement.

| $V_HH$-Fc | Wuhan-Hu-1 | D614G | N501Y | B.1.1.7 | B.1.351 | P.1 | B.1.617.2 |
|---|---|---|---|---|---|---|---|
| | $K_D$ (nM) | | | | | | |
| 8A2 | 0.26 | 0.2 | 0.11 | 0.001 | 0.001 | 0.5 | NB |
| 1B5 | 0.1 | 0.2 | 0.12 | 0.03 | 0.26 | 0.5 | 0.14 |
| 7A3 | 0.2 | 0.23 | 0.25 | 0.2 | 0.1 | 0.8 | 0.42 |
| 2F7 | 0.85 | 1 | 0.85 | 3.2 | NB | NB | NB |
| 8A4 | 0.16 | 0.6 | 0.5 | 5.6 | NB | NB | NB |
| 1H6 | 37 | 23 | 47 | 17 | NB | NB | NB |

NB: no binding

**Table S1.**

Affinity binding of SARS-CoV-2 $V_HH$-Fc fusions against the RBD of the original and variants on Octet.

| Assay | LP assay | PP entry assay | | | | Fluorescent reporter assay | | | | |
|---|---|---|---|---|---|---|---|---|---|---|
| $V_HH$ | Wuhan-Hu-1 | Wuhan-Hu-1 | B.1.1.7 | B.1.351 | P.1 | Wuhan-Hu-1 | D614G | B.1.1.7 | B.1.351 | P.1 |
| | $IC_{50}$ | $IC_{50}$ | $IC_{50}$ | $IC_{50}$ | $IC_{50}$ | $IC_{50}$ | $IC_{50}$ | $IC_{50}$ | $IC_{50}$ | $IC_{50}$ |
| | (nM) | (nM) | (nM) | (nM) | (nM) | (nM) | (nM) | (nM) | (nM) | (nM) |
| 8A2 | 5 | 14 | 2 | 1 | 0.5 | 74.8 | 17 | | | |
| 1B5 | 7.6 | 25 | | 5 | 5 | 127 | 22.2 | | | |
| 7A3 | 6.6 | 3 | 0.6 | 1 | 2 | 246 | 40.4 | | | |
| 2F7 | 8.6 | 15 | 10 | | | 120 | 26 | | | |
| 8A4 | 27.4 | | | | | 1388 | 160 | | | |
| 1H6 | 111.3 | | | | | | 308 | | | |
| 8A2+7A3 | 1.6 | 1 | 0.4 | 0.3 | 0.2 | 2.3 | 13.9 | 5 | 1.5 | 0.13 |
| 8A2+1B5 | 1.8 | 5 | 0.9 | 0.5 | 0.3 | 12.6 | 14.8 | 2.5 | 1.7 | 0.94 |
| 7A3+2F7 | 2.9 | 1 | 0.5 | 1 | 5 | 20.1 | 17.6 | 2.5 | 25.6 | 3.8 |
| 7A3+8A4 | 19.8 | | | | | 35.7 | | | | |
| 8A2+2F7 | 3.1 | | | | | 40.3 | 39.8 | 40 | 2.6 | 1.6 |
| 1B5+2F7 | 4.5 | 4 | 1 | 7 | 2 | 33.5 | 28.7 | 8 | 14.9 | 26.1 |

LP: lentivirus-based pseudovirus
PP: Pseudotyped particle

**Table S2.** Testing of SARS-CoV-2 $V_HH$-Fc fusion proteins and combination against the infection of pseudoviruses expressing the original and variant spike proteins.

| Assay | Live viral microneutralization assay | | | | | | Cytopathic assay |
|---|---|---|---|---|---|---|---|
| $V_HH$ | Wuhan-Hu-1 | D614G | B.1.1.7 | B.1.351 | P.1 | B.1.617.2 | Wuhan-Hu-1 |
| | $IC_{50}$ (nM) | | | | | | $IC_{50}$ (nM) |
| 8A2 | 72 | 23 | 17 | 2 | 0.16 | NB | 72 |
| 1B5 | 18 | 12 | 140 | 22 | 1 | | 242 |
| 7A3 | 46 | 4 | 25 | 7 | 1.4 | 19 | 56 |
| 2F7 | 36 | 13 | 186 | NB | NB | | 169 |
| 8A4 | | | | | | | 108 |
| 1H6 | | | | | | | 1125 |
| 8A2+ 7A3 | 1 | 6 | 2 | 0.87 | 0.14 | 27 | 20 |
| 8A2+ 1B5 | | | | | | | 47 |
| 7A3+ 2F7 | 1.5 | 12 | 5 | 27 | 1 | | 16 |
| 7A3+ 8A4 | | | | | | | 35 |
| 8A2+ 2F7 | 15 | 76 | 44 | 3 | 0.6 | | 97 |
| 1B5+ 2F7 | | | | | | | 40 |

**Table S3.**
Testing of SARS-CoV-2 $V_HH$-hFc fusion proteins and combination against Wuhan-Hu-1 and variant live viruses.

| Dataset | Hexapro/7A3 | Hexapro/8A2 | Hexapro/7A3/8A2 I | Hexapro/7A3/8A2 II |
|---|---|---|---|---|
| Microscope | Arctica | Arctica | Arctica | Krios |
| Voltage (KeV) | 200 | 200 | 200 | 300 |
| Detector | K2 | K2 | K2 | K3 |
| GIF slit (eV) | N/A | N/A | N/A | 20 |
| Frames | 60 | 60 | 60 | 50 |
| Exposure (s) | 9 | 9 | 7.8 | 4.49 |
| Flux (e/A) | 6.51 | 6.18 | 6.4 | 3.75 |
| Pixel size | 0.932 | 0.932 | 0.932 | 0.528 |
| Mode | Counting | Counting | Counting | Super-Res |
| Defocus Range | (-1.5, -2.2) | (-1.5, -2.2) | (-1.5, -2.2) | (-1.2, -2.2) |

**Table S4.** Cryo-EM datasets
